# Electrical stimulation of the human pulvinar generates visual percepts

**DOI:** 10.64898/2026.09.28.754866

**Authors:** William S. Kemball-Cook, Edward R. Bader, Alexis D. Boro, Emad N. Eskandar, Nathaniel J. Killian

## Abstract

Visual prostheses have targeted early structures in the visual pathway, including the retina, optic nerve, lateral geniculate nucleus, and visual cortex, to generate visual percepts. Stimulation of higher-order visual regions, which may better encode complex visual features, remains relatively unexplored. We tested whether electrical stimulation of the medial and inferior pulvinar elicits visual percepts in a patient with medically refractory epilepsy undergoing invasive seizure monitoring with intracranial electroencephalography. Bipolar stimulation (30 biphasic pulses, 0.5 ms per phase, 3 mA, 1 Hz) was delivered between adjacent pulvinar contacts while eye movements were tracked. Stimulation of different electrode pairs elicited visual percepts at distinct retinotopic locations (p = 0.010). Stimulation also altered cortical activity, producing an early power increase followed by gamma desynchronization in occipital and parietal regions (p < 0.001) and beta desynchronization in the lateral temporal region (p < 0.01). These findings suggest that pulvinar stimulation can generate visual percepts and modulate distributed cortical networks, supporting further investigation of the pulvinar as a target for vision restoration.

## Main Text

Millions of people worldwide suffer from irreversible vision loss [1]. One promising therapeutic approach for restoring vision is electrical stimulation of the visual pathway to generate visual percepts—phosphenes—that form the basis of artificial vision [2]. Visual prosthesis development has targeted early structures in the visual pathway, including the retina, optic nerve, lateral geniculate nucleus (LGN), and visual cortex [2]. In contrast, less work has examined stimulation of higher-order visual processing regions, which may offer advantages for encoding complex visual features [3]. Of particular interest is the pulvinar nucleus of the thalamus, which maintains extensive reciprocal connections with multiple visual cortical areas and serves critical roles in both visual attention and multisensory integration [4]. Previous reports in humans have provided mixed results regarding whether pulvinar stimulation generates visual percepts [5], and systematic mapping of percepts generated by pulvinar stimulation has not been attempted in humans.

Here, we examined whether electrical stimulation of the medial pulvinar (PuM) and inferior pulvinar (PuI) elicits phosphenes. This was tested in a single patient with medically refractory epilepsy due to extensive bilateral perisylvian and parieto-occipital polymicrogyria, who was undergoing invasive seizure monitoring with intracranial electroencephalography (iEEG).

Bipolar symmetric stimulation was delivered during 30-s epochs, each consisting of 30 individual biphasic pulses (0.5 ms per phase, 3 mA) at 1 Hz between adjacent contacts targeting the pulvinar. Prior to each stimulation epoch, the patient fixated on a crosshair at the center of an otherwise blank laptop screen. During stimulation, the patient was instructed to look at any visual sensations that appeared, verbally describe their characteristics, namely color, and indicate the approximate visual field zone in which they occurred by pressing a number (1–9) on a keypad (Fig. 1a). Eye-tracking data were collected throughout to monitor eye movements directed toward reported visual sensations. Additional details on eye-tracking, saccade detection, electrode localization, and iEEG data analyses are provided in the Supplementary Material.

**Fig. 1.**
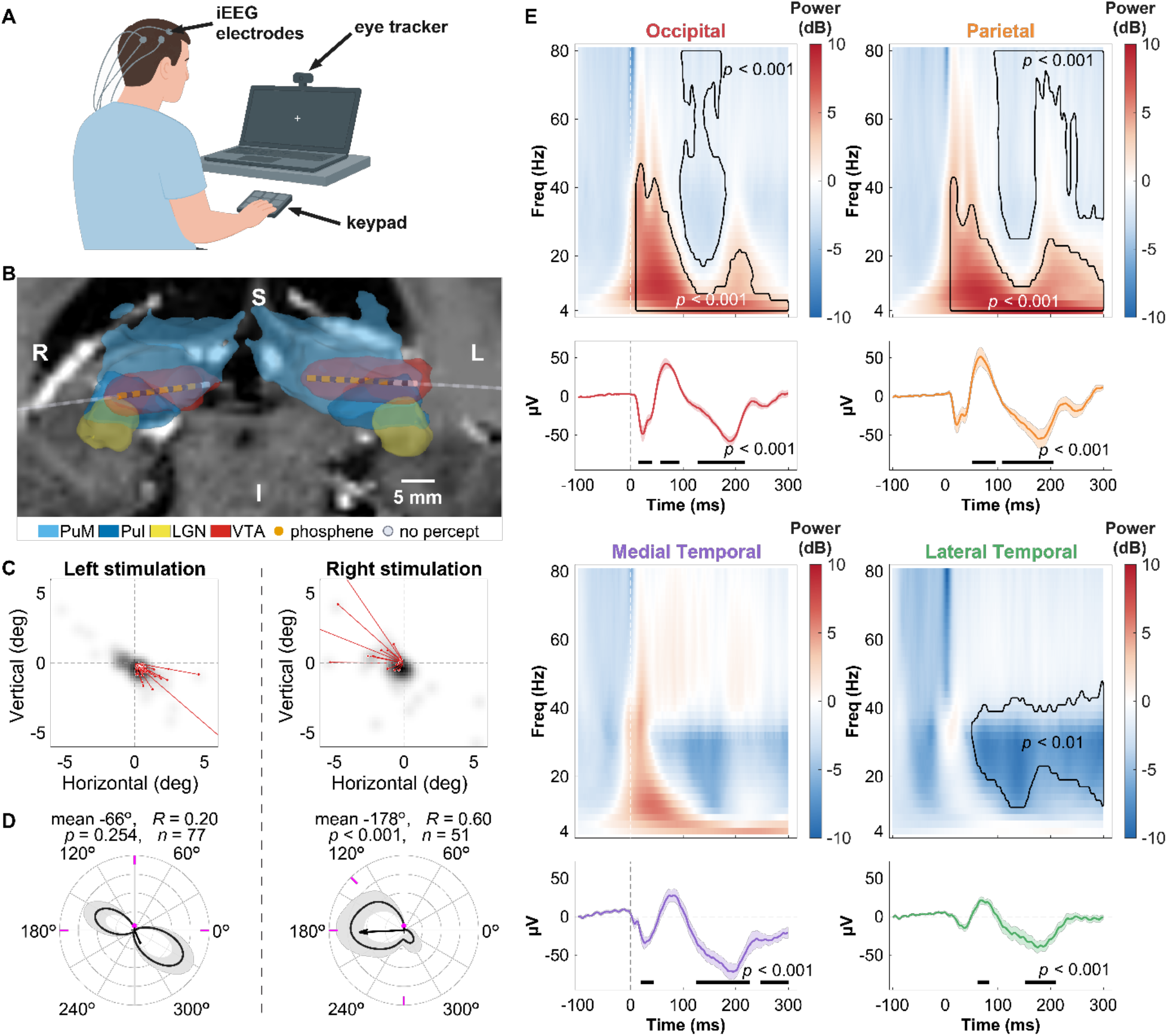
Behavioral and electrophysiological mapping of pulvinar stimulation. (a) Experimental apparatus: Before each stimulation epoch, the patient fixated a crosshair on a blank laptop screen. The patient reported all visual sensations, made saccades toward them, and indicated percept locations in the visual field via pressing keys on a number pad. (b) Electrode lead reconstructions; medial pulvinar (PuM, light blue), inferior pulvinar (PuI, dark blue), and lateral geniculate nucleus (LGN, yellow), with the union of the volume of tissue activated (VTA) from the 13 analyzed stimulation pairs in red. Phosphene-evoking contacts are shown in orange, and sites not involved in any reported percepts are shown in gray. (c) Aggregate phosphene location maps for stimulation of the left and right leads. Red lines show the individual saccade vectors, aligned to the origin, for a representative stimulation pair (left stimulation: contacts 5–6; right stimulation: contacts 3–4). Background maps show the fixation density pooled over all saccades directed toward percepts. (d) Circular density plots of saccades to percepts. Left panel: left pulvinar stimulation; right panel: right pulvinar stimulation. Kernel-density estimates are shown with 95% bootstrap confidence intervals. Magenta tick marks indicate the keypress directions. Directional clustering was assessed with a permutation test on the resultant vector length (left: R = 0.20, p = 0.254; right: R = 0.60, p < 0.001). Central percepts (keypress 5) were excluded from directional analyses. (e) Time-frequency representations (TFRs) of power (dB) and event-related potentials (ERPs, mean ± SEM) were normalized to a pre-stimulus baseline (−300 to −100 ms, dB) and are shown relative to stimulation onset for the cortical regions ipsilateral to stimulation. Outlines denote significant time-frequency clusters including low-frequency power increases and gamma band power decreases in the occipital and parietal regions (p < 0.001); a sustained beta desynchronization was observed in the lateral temporal region (p < 0.01). Bars on ERP panels mark significant time ranges (cluster-based permutation test, p < 0.001).

Electrode contacts in each hemisphere were localized to the PuM, PuI, and bordering the LGN (Fig. 1b). The patient reported visual percepts during stimulation epochs and reported distinct colors and locations for each percept (Table S1). Stimulation of different electrode pairs elicited phosphenes at distinct retinotopic locations, as indicated by saccades toward different visual field locations (permutation test, p = 0.010; Fig. 1c; see Supplementary Material). Right pulvinar stimulation resulted in saccades to percepts toward the contralateral hemifield (mean direction −178°, R = 0.60, p < 0.001, rotation permutation test; see Fig. 1d, Table S1). Saccade directions to percepts from left pulvinar stimulation were bilaterally distributed and, collectively, were not directionally clustered (mean direction −66°, R = 0.20, p = 0.254; see Fig. 1d, Table S1). These results suggest that pulvinar stimulation can evoke phosphenes predominantly in the contralateral hemifield, but with potentially ipsilateral visual field encoding as well.

We computed time-frequency power relative to stimulus onset to understand the effects of pulvinar stimulation on cortical activity (Fig. 1e). Pulvinar stimulation resulted in an immediate and prolonged power increase (cluster median 12 Hz, 4 to 46 Hz) and a later gamma band desynchronization in the occipital and parietal regions (cluster-based permutation, p < 0.001). In contrast to the posterior gamma effect, stimulation produced an early, prolonged beta-range desynchronization in the lateral temporal region (median 30 Hz, from 51 ms, p < 0.01); beta suppression (14–30 Hz, 100–300 ms) was greater in the medial and lateral temporal regions than in the occipital and parietal regions (Wilcoxon rank-sum, z = −2.63, p = 0.009). These findings suggest that pulvinar stimulation engages a distributed cortical network that may support the encoding of visual percepts.

To our knowledge, this is the first demonstration in humans that pulvinar stimulation produces visual percepts. Saccades were typically directed toward the contralateral hemifield as expected [6]. Interestingly, however, some sites evoked percepts in the ipsilateral hemifield. Past pulvinar studies have focused predominantly on the inferior and lateral pulvinar, termed the “visual pulvinar”, which contain two retinotopic maps of contralateral visual space in non-human primates [7]. The PuM has received comparatively less attention and is considered a multimodal integration center given its broad cortical connectivity [4]. However, several studies have identified regions within the PuM that show weak contralateral tuning, suggesting it contains large receptive fields encompassing ipsilateral space [7, 8]. Thus, the PuM itself may contribute through its dense projections to visual cortical areas.

Pulvinar stimulation modulated activity across the occipital, parietal, and temporal regions. Prior studies have demonstrated anatomic and functional connectivity between the pulvinar and these regions in both humans and non-human primates [4, 9, 10]. In our study, the engagement of the parietal and temporal regions following pulvinar stimulation is consistent with this connectivity. Moreover, these results suggest that phosphene perception could be partially mediated by direct pulvinar projections to higher-order cortical areas rather than solely direct visual cortex activation. We found that pulvinar stimulation suppressed beta-range activity in the temporal lobe, which may reflect differential engagement of pulvinar subregions, as prior studies have demonstrated that the dorsal and ventral PuM preferentially connect to the parietal and temporal lobes, respectively [7].

This study provides evidence that electrical stimulation of the pulvinar can create visual percepts and modulate visual cortical networks, thus supporting the pulvinar as a target region for vision restoration. Furthermore, modulation of cortical regions that contribute to object recognition, visual memory, and spatial attention may offer advantages for visual prostheses. Targeting such regions alongside early visual areas could support richer, more naturalistic percepts, and stimulation paradigms could be developed to help engage attention to relevant stimuli, such as low-lying objects or navigational hazards. However, future studies should reproduce these findings in additional subjects and employ additional controls such as sham stimulation. New technologies that enable higher-density bilateral cortical sampling could improve pulvinar stimulation-based connectomics and interrogate the widespread effects on cortical activity that we observed.

