## Supplementary Material for "Electrical stimulation of the human pulvinar generates visual percepts"

### Supplementary Materials

*Ethics Statement*

Electrical stimulation of the pulvinar as reported in this study was performed as part of the patient's standard clinical care during stereoelectroencephalography for drug-resistant epilepsy, and not as a separate research procedure. The pulvinar nucleus is recognized as a target for chronic stimulation to reduce seizure frequency, including with responsive neurostimulator implants (NeuroPace, Inc.). Implantation of neurostimulation devices targeting the pulvinar is often performed in patients who are not candidates for surgical resection of the seizure focus and who have drug-resistant epilepsy with seizures arising in the posterior quadrant, posterior lateral temporal areas, or are multifocal [1, 2]. In patients being evaluated as potential candidates for pulvinar neurostimulation, direct electrical stimulation of the pulvinar during intracranial monitoring is used as part of standard clinical care to confirm the feasibility of stimulation, characterize stimulation-evoked responses, and screen for perceptual or other adverse effects prior to possible device implantation [3, 4]. Accordingly, in keeping with Albert Einstein College of Medicine Institutional Review Board policies, ethical approval was not required for acquisition of the data described in this study. The data described herein were acquired in accordance with the Declaration of Helsinki. Written informed consent from the patient was acquired for publication of this case report. During stimulation, no discomfort was reported and no seizures occurred. The patient was subsequently implanted with a responsive neurostimulation system, with one occipital lead and one pulvinar lead, and has reported no adverse events related to the implant or its use.

*Eye-tracking and Saccade Detection*

Eye-tracking was performed using a USB camera (120 fps, 720p resolution) mounted on a Lenovo ThinkPad P51 laptop with a 15.6"-diagonal display. We created custom C++ software for this study using the Eyedid SDK. The software sent a per-frame TTL synchronization pulse to the Natus Quantum amplifier used for post hoc alignment. The subject sat approximately 40 cm from the screen, which subtended approximately 48 × 28 degrees of visual angle (DVA). Before the session started, gaze position was calibrated using a homography transform. Gaze position was recentered before each stimulation at each site pair by capturing the fixation of a central point. Tracking gaps longer than 100 ms were excluded from analysis. Following our previously used saccade detection parameters [5], gaze position was low-pass filtered (30 Hz cutoff) and a saccade speed threshold of 25 degrees per second was used in the 40–500 msec post-pulse windows. To avoid counting saccades toward the keypad and saccades back toward the screen, any saccades occurring within 500 ms of a keypress were excluded, and for each stimulation site, saccade directions were clustered using circular k-means (k = 3), and the cluster with at least 33% of the saccades and with the largest resultant vector was retained. A vector from the start to end gaze locations defined each saccade, and these vectors were realigned to the origin for subsequent analysis, i.e., assuming retinocentric percepts.

Retinotopic specificity was tested by comparing the spatial spread of per-pair mean saccade displacements against a null distribution obtained by shuffling the bipolar stimulation contact pair label for each saccade (10,000 permutations). Directional clustering within each stimulated hemisphere was assessed by rotating each stimulation pulse's saccades by a random angle and recomputing the resultant vector length (2,000 permutations).

*Electrode Localization*

Depth electrodes from PMT Corporation were used (16-contact, 1.3-mm contact length, 0.8-mm diameter, 2.2-mm spacing) targeting the pulvinar. Remaining leads ranged in configuration from 12–16 contacts with 1.3–2 mm contact lengths and 2.2–4.43 mm spacing. The pre-operative T1-weighted MRI was processed using the FreeSurfer automated segmentation and surface reconstruction pipeline [6]. Post-operative CT scans were co-registered to the T1-weighted MRI, and electrode coordinates were identified using the MATLAB FieldTrip Toolbox [7].

Pulvinar stimulation contacts were assigned to the medial (PuM) and inferior (PuI) pulvinar based on the Morel atlas [8]. The most ventral pulvinar contacts bordered the lateral geniculate nucleus (LGN). Remaining contacts were localized using the Desikan-Killiany cortical parcellation [9] and FreeSurfer subcortical segmentation. White matter contacts were excluded from the analysis. Based on the subject's distribution of electrodes, contacts were grouped into four macro-regions: medial temporal lobe, lateral temporal lobe, occipital lobe, and parietal lobe.

*iEEG data processing*

iEEG signals were recorded using Natus software (Natus Medical Incorporated, Middleton, WI) at 2048 Hz. Stimulation artifacts were removed using the signal replacement method described in Crowther et al [10]. For each stimulation pulse, a 10 ms artifact window was extracted. Pre- and post-artifact segments (5 ms each) were time-reversed, resampled to the artifact window duration, linearly tapered, and summed to replace the artifact.

Re-referencing was performed using an independent component analysis (ICA) approach described in Michelmann et al [11]. First, large artifacts were identified as amplitude deviations exceeding 5 interquartile ranges above the median on three or more channels within 300 ms windows. Data were then band-stop filtered (55–65 Hz, 115–125 Hz, 175–185 Hz) to remove line noise, and ICA was computed on the artifact-free data. The recording was subsequently interpolated using Piecewise Cubic Hermite Interpolating Polynomial interpolation in 150 ms windows around moments where artifacts had been marked. The three components with the highest spatial broadness scores were removed prior to signal reconstruction.

Local field potentials (LFPs) were first processed with discrete Fourier transform regression line noise removal at 0.1 Hz resolution and were then zero-phase bandpass filtered (1–200 Hz for time-frequency analyses and 1–100 Hz for ERPs, 2nd-order Butterworth). Analysis epochs were defined as −700 to +700 ms from each stimulation pulse onset. Time-frequency power was computed using a Morlet wavelet method (4 to 80 Hz, 3 cycles) and normalized to the pre-stimulus baseline (−300 to −100 ms). Power was averaged across the stimulation events for each channel and then averaged across channels within each larger brain region. Left and right hemisphere stimulation epochs were analyzed separately using only the ipsilateral hemisphere contacts, and then pooled (Fig. 1e). Stimulation-evoked time-frequency changes in 10–300 msec post-stimulus were tested against the pre-pulse baseline using two-sided cluster-based permutation tests (dependent-samples t statistic, 10,000 permutations, cluster-forming $\alpha = 0.01$, cluster-level $\alpha = 0.025$) with the first 10 ms after each pulse excluded to avoid residual stimulation artifact.

*Volume of tissue activated (VTA) modeling*

The volume of tissue activated (VTA) by each bipolar stimulation pair was estimated by finite-element modeling using OSS-DBS [12]. The electrodes were modeled based on manufacturer specifications (PMT Corporation): 16 contacts, 1.3-mm contact length, 2.2-mm pitch, and 0.8-mm diameter. We modeled a 0.1-mm encapsulation layer, and dielectric properties were simulated with four Cole-Cole dispersions [13] with gray matter, white matter, and CSF maps segmented from the T1-weighted MRI. White-matter conductivity anisotropy was mapped from the diffusion tensors [14]. The electric fields from the ±3-mA pulses were computed and evaluated at 1 kHz on a 48 × 48 × 48 lattice with 0.3-mm resolution, centered between the two active contacts. The VTA was defined as the region where the field exceeded 0.103 V/mm, calculated by scaling a 0.2 V/mm threshold for a 90-µs pulse width [15] to the 0.5-ms pulse width, assuming a 129-µs chronaxie for the thalamus [16]. The VTA volumes were pooled across the 13 analyzed stimulation pairs (1028 $mm^3$ union) for display in Fig. 1b. VTA-nucleus overlap is indicated, per pair, in Table S1.

| Stim Contacts | Nearest Nucleus | PuM VTA (%) | PuI VTA (%) | LGN VTA (%) | Saccades (n) | Direction (°) | Eccentricity (°) | Latency (ms) | Phosphene Color | Keypress |
|---|---|---|---|---|---|---|---|---|---|---|
| 1–2 | Right PuM | 1 | 0 | 0 | - | - | - | - | None | - |
| 2–3 | Right PuM | 53 | 0 | 0 | 26 | +179.1 ± 6.5 | 4.99 ± 2.13 | 264.4 ± 27.8 | White | 7 |
| 3–4 | Right PuM | 70 | 7 | 0 | 17 | +163.8 ± 7.8 | 3.62 ± 1.45 | 274.8 ± 35.3 | Black | 4 |
| 4–5 | Right PuM | 69 | 20 | 0 | - | - | - | - | None | - |
| 5–6 | Right PuI | 57 | 38 | 2 | 24 | -75.8 ± 5.9 | 1.29 ± 0.31 | 308.2 ± 24.5 | White | 5 |
| 6–7 | Right PuI | 32 | 46 | 11 | 8 | -49.4 ± 7.7 | 3.37 ± 1.71 | 294.2 ± 40.3 | Yellow | 2 |
| 1–2 | Left PuM | 97 | 0 | 0 | 31 | -55.2 ± 5.1 | 1.31 ± 0.31 | 253.9 ± 27.1 | White | 5 |
| 2–3 | Left PuM | 99 | 0 | 0 | - | - | - | - | None | - |
| 3–4 | Left PuM | 96 | 3 | 0 | 32 | +159.7 ± 2.9 | 2.18 ± 0.70 | 243.1 ± 25.2 | Yellow | 4 |
| 4–5 | Left PuM | 84 | 12 | 0 | 0 | - | - | - | Yellow | 6 |
| 5–6 | Left PuM | 70 | 21 | 0 | 45 | -34.9 ± 2.7 | 1.29 ± 0.27 | 296.3 ± 19.5 | White | 5, 8 |
| 6–7 | Left PuM | 52 | 28 | 1 | - | - | - | - | None | - |
| 7–8 | Left PuI | 33 | 26 | 2 | - | - | - | - | None | - |

**Table S1. Characteristics of saccades made to locations of phosphenes produced during visual thalamus stimulation epochs.** Bipolar stimulation was delivered between adjacent contacts in 30-second epochs. The nearest visual thalamic nucleus is indicated, and the PuM, PuI, and LGN columns report the percent of the estimated VTA within that nucleus (Fig. 1b). Saccade count is given, and saccade direction (0° = rightward, increasing counterclockwise), eccentricity, and latency are reported as mean ± SEM. The patient self-reported phosphene color (stimulation sites without a reported percept are indicated as 'None'). The percept location was indicated via a numeric keypad: the Keypress column lists the reported keypad digit (1 to 9; 5 = center; keypad grid 7-8-9 / 4-5-6 / 1-2-3). For directional analyses and Fig. 1d, central percepts (keypress 5 only) were excluded, giving 77 saccades for left stimulation and 51 saccades for right stimulation.

| **Cortical Region** | **Left Hemisphere** | **Right Hemisphere** | **Total** |
|---|---|---|---|
| Medial Temporal | 3 | 12 | 15 |
| Lateral Temporal | 4 | 12 | 16 |
| Occipital | 4 | 15 | 19 |
| Parietal | 1 | 12 | 13 |
| **Total** | **12** | **51** | **63** |

**Table S2.** Distribution of electrodes across cortical regions. Electrode coordinates were localized to cortical regions using the Desikan-Killiany cortical parcellation and FreeSurfer subcortical segmentation.

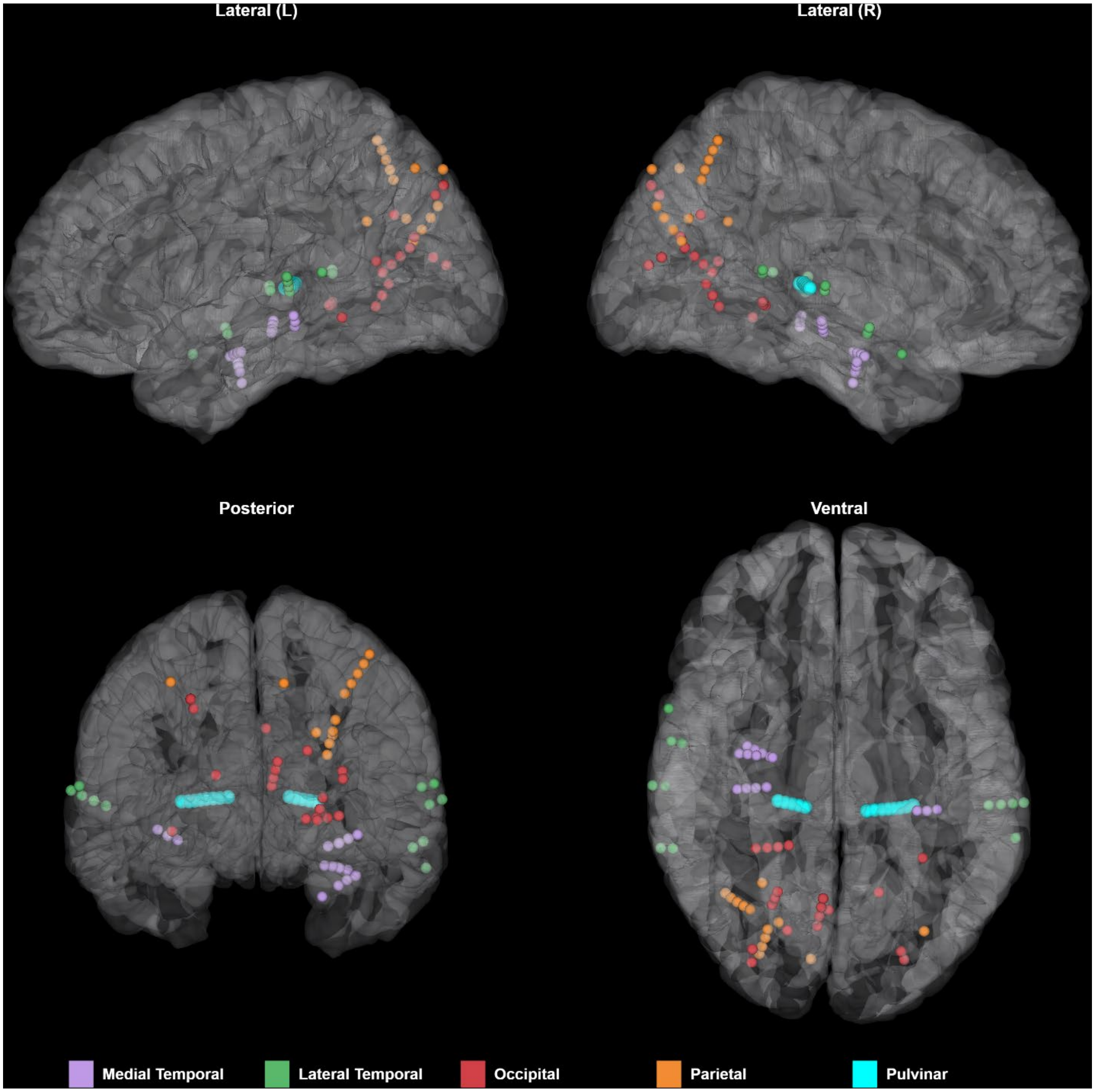


**Figure S1.** Electrode locations projected onto the Colin27 FreeSurfer pial surface. Recording contacts are color-coded by macro-anatomical region. Pulvinar stimulation contacts are shown in cyan. Contacts outside of the pial surface are not shown.
